# Symbiosis with *Rhizophagus irregularis* improves grapevine rootstock performances under water deficit conditions

**DOI:** 10.64898/2025.12.17.694802

**Authors:** Lucas Galimand, Martin Lamy, Laure Valat, Hélène Laloue, Yann Leva, Laurence Deglène-Benbrahim, Loïc Yung, Julie Chong

## Abstract

Grapevine represents a major crop with crucial socio-economic importance, however, its culture is threatened by climate change, especially drought. Indeed, water deficit has a negative impact on grapevine growth and yield but also impacts fruit and wine quality. To improve grapevine resilience to drought, developing strategies such as symbiosis with Arbuscular Mycorrhizal Fungi could be a promising. We focused on the benefit of using *Rhizophagus irregularis* DAOM 197198 for improving performances in controlled conditions of two highly used rootstocks (41B and SO4) under moderate to severe water deficit. At a field capacity of 14-40%, SO4 was more affected compared to 41B. Successful functional symbiosis was obtained for the two rootstocks, both in well-watered and water deficit conditions. Interestingly, colonization with *R. irregularis* improved growth and photosynthetic parameters in both 41B and SO4, especially under water stress, restoring them to the levels on non-stressed plants. Further analysis of mineral nutrition and aquaporin expression revealed contrasting responses between the two rootstocks. Whereas mycorrhization strongly enhanced phosphorus contents in both 41B and SO4 roots and leaves, the overall beneficial effects of the symbiosis on mineral nutrition were more pronounced in SO4. In contrast, the expression of *VvPIP2.1*, a highly water-permeable aquaporin involved in root hydraulic conductivity was increased in mycorrhized roots of 41B but repressed in SO4. This study emphasizes that interaction between AMF and grapevine induce contrasting effects on plant nutrition depending on the rootstock genotype, and that mycorrhizal inoculation could be of interest in the case of drought sensitive rootstocks.

## Introduction

Today, developing strategies for crop adaptation to changing environment is crucial. Climate change results in several biotic and abiotic stresses severely impacting yield and food security worldwide. Drought, driven by scarcity or absence of rainfalls and elevated temperatures, has become a critical concern regarding abiotic stresses limiting crop sustainability and productivity (Raza et al. 2019; Dietz et al. 2021).

Among crops of economic significance, grapevine is a perennial plant cultivated worldwide (7.3 Mha global cultivating area) with high economic importance. However, grapevine is threatened by climate change, especially in European vineyards (Wolkovich et al. 2025). Heat and drought stresses impact Mediterranean areas (Songy et al. 2019), thus water deficit becomes increasingly worrying in many traditional wine producing regions (Gambetta et al. 2020).

Water deficit has a negative impact on grapevine growth and yield but also impacts fruit and wine quality (Chaves et al. 2010). Water deficit alters grapevine growth through inhibition of the photosynthetic machinery (Ju et al. 2018; Cogato et al. 2022) and could further trigger embolism formation within the xylem, resulting in wine mortality (Leeuwen and Darriet 2016; Gambetta et al. 2020). In addition, water deficit alters the quantity and the quality of berries, leading to modification of fruit composition and wine quality (Chaves et al. 2010; Torres et al. 2018).

Grapevine is cultivated as grafted plant and its sensitivity to drought depends on both rootstock and scion. Rootstocks show contrasted tolerance to water depletion and play an important role in grapevine resilience to this stress (Tramontini et al. 2013). However, grapevine culture depends on the use of a very limited number of rootstock genotypes, 90% of *V. vinifera* scions being grafted onto less than 10 rootstock genotypes. Low genetic variability of rootstocks could thus limit the adaptation capacity of the plant (Bernardo et al. 2025). To cope with water deficit, grapevine is able to develop several adaptative responses such as stomata closure, which is regulated by the plant hormone abscisic acid, allowing transpiration reduction and avoiding critical water potential that could lead to hydraulic failure (Nakashima and Yamaguchi-Shinozaki 2013; Gambetta et al. 2020). Osmolyte accumulation, especially amino acids such as proline, calcium or other inorganic ions have been reported to play important role in leaf osmotic adjustment during water stress (Degu et al. 2019; Gambetta et al. 2020). Aquaporins (AQP) are also key players in adaptation of vines to drought conditions. Aquaporins are Major Intrinsic proteins, a large superfamily of conserved small membrane channels facilitating the transport of water and small neutral molecules (Maurel et al. 2015; Braidotti et al. 2024). AQP are involved in the cell-to-cell radial water transport from soil into root xylem vessels (Maurel et al. 2015) and at the plant level they are involved in the rapid and reversible regulation of cell hydraulic conductance (Sabir et al. 2021). In addition, AQP expression is modulated during water stress conditions in several plant species in accordance with a role as important players of water status regulation. Aquaporins appears as interesting candidates involved in grapevine adaptation to water deficit since modulation of their expression has been linked to effects on several hydraulic traits (Braidotti et al. 2024).

In order to improve grapevine resilience to drought, the use of beneficial microorganisms appears to be a promising strategy. Among beneficial microorganisms, Arbuscular Mycorrhizal Fungi (AMF) are obligate symbionts from the Glomeromycota phylum that contributes to several ecosystem services, by improving plant nutrition and immunity (Shi et al. 2023). Grapevine, like 80% of terrestrial plants, naturally establishes symbiosis with AMF. However, grapevine-AMF symbiosis dynamic and benefits rely on management system practices and rootstocks genotypes (Nikolaou et al. 2003; Moukarzel et al. 2022; Velaz et al. 2025). Several mechanisms could explain the beneficial effect of AMF in plant adaptation to drought: mycorrhizas, through hyphal network, allow roots to explore a larger volume of soil, giving access to a higher amount of water and nutrients. Plants colonized by AMF can also obtain their phosphorus through the mycorrhizal pathway, which frequently proves more efficient at mobilizing soil phosphorus than direct root uptake (Smith et al. 2011), and particularly in the context of water limitation (Püschel et al. 2021). Finally, AMF colonization is known to enhances photosynthesis performance and osmolyte accumulation (Trouvelot et al. 2015; Velaz et al. 2025).

Several studies described a positive impact of symbiosis with AMF on grapevine tolerance to water stress in both controlled conditions and in the vineyards. Cabernet Sauvignon grafted on drought sensitive rootstocks inoculated with *Funneliformis mosseae* showed significantly improved drought resistance as well as enhanced foliar growth and P concentration compared to non-inoculated plants (Nikolaou et al. 2003). In another study, Ye et al. (2023) further highlighted that a commercial mix of AMF could mitigate the effect of drought stress in *V. vinifera* cv. Ecolly by regulating the osmotic and antioxidant balance as well as the expression of drought responsive genes. In the same way, Kozikova et al. (2024) reported that a commercial inoculum containing a mixture of five AMFs and beneficial rhizobacteria improved the performance of Tempranillo and Cabernet Sauvignon grafted on 110-R under water deficit conditions. In field conditions, inoculation of AMF (*Funneliformis mosseae*) through rye donor plants had a positive impact on grapevine (Viosinho grafted onto 1103 Paulsen rootstock) photosynthesis and yield after a heat wave (Nogales et al. 2021). In addition, under exacerbated summer stress, Cardinale et al. (2022) reported a higher survival rate as well as enhanced element content in mycorrhized rootstock 1103-Paulsen compared to non-mycorrhized plants.

Despite the recent attention paid to the use of beneficial microorganisms, especially AMF to mitigate the effect of climate change in grapevine, there are few studies of the effect of the single strain *Rhizophagus irregularis* (Ri) on rootstock performance in water deficit conditions (Aguilera et al. 2022; Dagher et al. 2025). Indeed, most of studies using AMF for drought resilience focused on commercial mixtures of several AMF. *R. irregularis* DAOM 197198 is the most used species in commercial bioinoculants and has become the model species for AMF research (Kokkoris et al. 2024). In addition, with the growing prevalence of water stress in vineyards, it is essential to evaluate the potential benefits of this strain for the rootstocks used in increasingly vulnerable areas. In this context, this work aimed at evaluating the effects of *R. irregularis* DAOM 197198 on performances of two highly used rootstocks in French vineyards: 41B (*Vitis vinifera* X *Vitis berlandieri*) and SO4 (*Vitis riparia* X *Vitis berlandieri*), under water deficit. 41B and SO4 have been reported to have medium and low-medium tolerance to drought respectively (Lovisolo et al. 2016). We further characterized the responses of rootstocks to both AMF symbiosis and water deficit by studying leaf and root growth, photosynthesis parameters, as well as water and mineral nutrition with a special focus on root aquaporin expression.

## Materials and Methods

### 1. Plant cultivation and AMF inoculation

Plantlets of 41B MGt (*Vitis vinifera* cv. Chasselas × *V. berlandieri*) and SO4 (*V. berlandieri* × *V. riparia*) were initially propagated *in vitro* on Woody Plant Medium from single-node cuttings. After 6 weeks, plantlets were transferred into individual 1 L plastic pots filled with a sterilized sand and perlite mixture (1v/1v) for *ex vitro* acclimatization. The pots were then placed in a growth chamber with 16 h of light at 25 °C and 8 h of darkness at 19°C, 60% relative humidity and a light irradiance of 150 µE m^−2^ s^−1^. Plantlets were watered with a complete nutrient solution containing 0.5 mM KH_2_PO_4_ twice a week as described in (Valat et al. 2018). Four weeks after acclimatization, young vines were transferred to 1,5 L plastic pots filled with 200 g of sterile gravel in the bottom and 600 g of a sterile mix of sand and perlite (1v/1v). Plants were subsequently watered during one week with a low-Pi nutrient solution (0.1 mM KH_2_PO_4_).

One week after repotting, 20 plants of each genotype were individually inoculated with 1 000 spores of *Rhizophagus irregularis* DAOM 197198 (Ri condition), produced under axenic conditions (Agronutrition, Carbonne, France). AMF inoculation followed the protocol described in Goddard et al. (2021). Briefly, the inoculum was diluted with tap water to obtain a suspension of 50 spores per mL. Then 20 mL of this diluted inoculum were gently poured at the base of the stem. Non-inoculated plants (n = 20 for each genotype) were watered with 20 mL of tap water (NoRi condition). Plants were watered for one week with tap water to promote AMF symbiosis, then twice a week with the low-Pi nutrient solution for 6 weeks before water deficit implementation.

### 2. Hydromineral deficit treatments

Field capacity (FC) of the sand and perlite mix was evaluated prior to the implementation of drought stress. Five pots filled with this substrate were saturated with water (100% FC). Pots at saturation were weighed, then dried out at 105 °C for 24 h and weighed again to determine the water mass in pots at saturation. Ratio of water mass / dry substrate mass was equivalent to 0.18 (*i.e.* at 100 % FC, water content was equivalent to 18 % of the substrate dry weight). This calculated FC was used to determine the irrigation volumes required for each plant under each condition.

Hydromineral deficit treatments were initiated six weeks after the plant inoculation. At each watering, pots were individually weighed, and low-Pi nutrient solution was added to reach 40 % FC for water deficient condition (WD) and 100 % FC for control well-watered plants (NoWD). Between two waterings, stressed plants were maintained between 14 % and 40 % FC corresponding to moderate-to-severe water deficit. Well-watered plants were maintained between 65 % and 100 % FC, corresponding to hydric conditions favorable for vine growth. Due to the highly draining nature of the substrate, plants required irrigation three times per week to maintain the water conditions. Hydromineral stress was applied during 5 weeks before harvesting the plants. In total, the experiment comprised 80 plants (2 genotypes x 2 inoculation treatments x 2 water conditions x 10 replicates).

### 3. Assessment of mycorrhizal parameters

At the end of the experiment, root mycorrhizal parameters were assessed on 6 randomly selected plants for each modality. Root segments of selected plants were cleared in 10% (w/v) KOH solution with 30 % H_2_O_2_ solution and then immersed in an ink solution (5% Black Sheaffer Skrit ink / 8% acetic acid) to stain fungal structures, following the ink-based protocol described in Vierheilig et al. (2005). The samples were cut into 10-mm pieces and then observed under an optical microscope. The parameters related to the mycorrhizal colonization were calculated according to Trouvelot et al. (1986).

### 4. Growth and biomass measurements

Growth and biomass measurement were realized on the 10 plants of each modality. Plant height was measured using a measuring tape. Each plant produced 1 to 3 shoots, and growth was quantified as the difference in cumulative shoot height between the onset of hydromineral deficit and the end of the experiment. Five weeks after inoculation, shoots and roots of each plant were harvested, separated and cleared from culture substrate particles. Total fresh weight (FW) of each sample was determined immediately at harvest. After collecting various samples for subsequent analysis, remaining FW was weighed again. Remaining root and shoot samples were then oven-dried at 70 °C for 48 h to determine the dry weight (DW).

### 5. Analysis of trace and major elements in plants

The root and leaf elemental content were determined for the 6 plants previously selected for mycorrhizal colonization assessment. Dried leaves and roots were transferred into 20 mL polyethylene oscillation vials (FisherbrandTM, Thermo Fischer Scientific, USA) and individually ground to a fine homogeneous powder using a beadmill (MM400, Retsch, Germany).

The dried powder was digested according to the protocol detailed in Yung et al. (2021), with either 50 mg of root powder or 500 mg of leaf powder. The quantities of H_2_O_2_ and HNO_3_ were calibrated in accordance with the mass of the samples. Trace (i.e. Fe, Mn and Zn) and major (i.e., Ca, K, Mg, K, P, and S) elements were analyzed by inductively coupled plasma optical emission spectrometry (ICP-OES, iCap 600, Thermo Fischer Scientific, Inc., Pittsburgh, USA). To ensure the accuracy and precision of the analytical process, control samples with known elemental compositions (oriental basma tobacco leaves certified material, INCT-OBTL-5, LGC Promochem, Molsheim, France) were included in the analyses.

Whole plant element contents were calculated by adding leaf and root values for each replicate.

### 6. Photosynthesis measurements

Four weeks after the implementation of the hydromineral deficit treatment, chlorophyll fluorescence and gas exchanges were measured on intact leaves of the 6 previously selected plants using a LI-6800 portable photosynthesis system (LI-COR Biosciences, USA). Measurements were conducted on three fully-expanded leaves, starting from the apex, for each plant. Gas exchanges parameters included net CO_2_ assimilation rate (A, µmol CO_2_.m^-2^.s^-1^) and stomatal conductance (gsw, mmol H_2_O.m^-2^.s^-1^). For each leaf, 30 measurements were recorded at 6 s intervals during daytime under controlled conditions: air flow 600 µmol.s^-1^, CO_2_ concentration 450 µmol.mol^-1^, photosynthetic photon flux density 200 µmol m^-2^.s^-1^, leaf temperature 25°C and 55% relative humidity in the measuring chamber. Fluorescence parameters included maximum photochemical efficiency of PSII (Fv/Fm) and quantum yield of PSII (ΦPSII). Fluorescence was measured in the dark after overnight dark adaptation to ensure that all photosystem II reaction centers were open before being saturated by light.

### 7. Aquaporins gene expression analysis using RT-qPCR

Immediately after harvesting, 65-75 mg of roots were sampled, flash-frozen in liquid nitrogen, finely ground using a mill, and stored at -80°C. RNA extractions were performed as described in Duret et al. (2022) first using ConcertTM Plant RNA Reagent (InVitrogen) after addition of PVP-40 (3% m/v final). RNA was subsequently extracted with Direct-zolTM RNA kit (Zymo Research, Irvine, CA, USA). DNase I treatment was realized during RNA extraction following manufacturer’s protocol. RNA quantification and purity were assessed with an optical density control (NanoDrop One^TM^, Thermo Fischer Scientific, USA) based on OD260/280 and OD260/230 ratios. Integrity of RNA and absence of genomic DNA was checked by 1% agarose gel electrophoresis. Each cDNA was converted from 500 ng of DNA-free RNA using the iScriptTM Reverse Transcription Supermix® for RT-qPCR (Bio-Rad, USA) following the manufacturer’s instructions. Quantitative PCR was performed with a CFX Duet Real-Time PCR detections System (Bio-Rad, USA) using iTaq Universal SYBR® Green Supermix (Bio-Rad, USA). Each 25 μL reaction contained 12.5 μL of SYBR Green Supermix, 0.2 μM of forward and reverse primers, and cDNA corresponding to 10 ng of reverse-transcribed RNA. The thermal cycling program was: initial denaturation at 95 °C for 3 min, followed by 40 cycles of 95 °C for 15 s, 60 °C for 20 s, 72 °C for 20 s, 76°C for 20 s (plate read). Melting curve analysis (60–95 °C, increments of 0.5 °C every 5 s) were used to verify amplification specificity. Nine genes encoding grapevine aquaporins representative from different families were selected as targets (Tab. S1): *VvNIP5.1*, *VvNIP1.1* (nodulin26-like intrinsic proteins, NIPs), *VvPIP2.1*, *VvPIP2.2*, *VvPIP1.2*, *VvPIP1.4* (plasma membrane intrinsic proteins, PIPs), *VvTIP1.1* and *VvTIP 2.2* (tonoplast intrinsic proteins, TIPs) and VvXIP (uncharacterized intrinsic proteins, XIPs). *GintPT* was analyzed a as marker of symbiosis functionality (Duret et al., 2022). Two reference genes, *VvEF1a* and *VvActin* were used for normalization. Relative expression levels were calculated using the 2^−ΔΔCt^ method (Schmittgen and Livak 2008). Well-watered non-mycorrhizal plants (NoRi – NoWD) of the same genotype were used as the calibrator. Results are analyzed and represented as the log_2_ of the raw data.

### 8. Statistical analysis

All statistical analyses were performed with RStudio v.4.4.2. Data sets were first analyzed using descriptive statistics for each variable. For each genotype a linear model was then constructed including the treatment factors (*R. irregularis* inoculation and water regime). Normality of model residuals and homoscedasticity of variances were assessed using respectively the Shapiro–Wilcoxon test and Levene’s test. When both assumptions were verified, data were analyzed by one-way ANOVA. Post-hoc comparisons were conducted using Tukey’s HSD test. When normality or homoscedasticity of data could not be checked, a non-parametric Kruskal–Wallis test was applied with Dunn’s post-hoc test and Holm correction. Differences were considered statistically significant at *p* < 0.05.

## Results

### 1. Successful symbiosis between *Rhizophagus irregularis* and both 41B and SO4 rootstocks genotypes

No fungal structure was observed in fragments without *R. irregularis* inoculation (NoRi) in both 41B and SO4 genotypes (Fig. 1B). Regarding inoculated plants, both genotypes showed high frequency rates of mycorrhizal colonization reaching F% = 95-100 % in both NoWD and WD conditions (Tab. 1). In SO4, plants exposed to drought stress showed significant lower intensity of root colonization and arbuscule abundance. In contrast, for 41B genotype, no significant difference between NoWD and WD condition was observed concerning M (%) and A(%) parameters, suggesting no effect of the water regime on intensity and functionality of mycorrhizal symbiosis. High expression of GintPT, a marker of symbiosis functionality (Duret et al. 2022) was observed in both rootstocks in well-watered as well as in water deficit conditions (Fig. 1A). These results confirmed the success of mycorrhization with *R. irregularis* and show that symbiosis is well established in plants under both NoWD and WD conditions (Fig. 1C).

**Figure 1.**
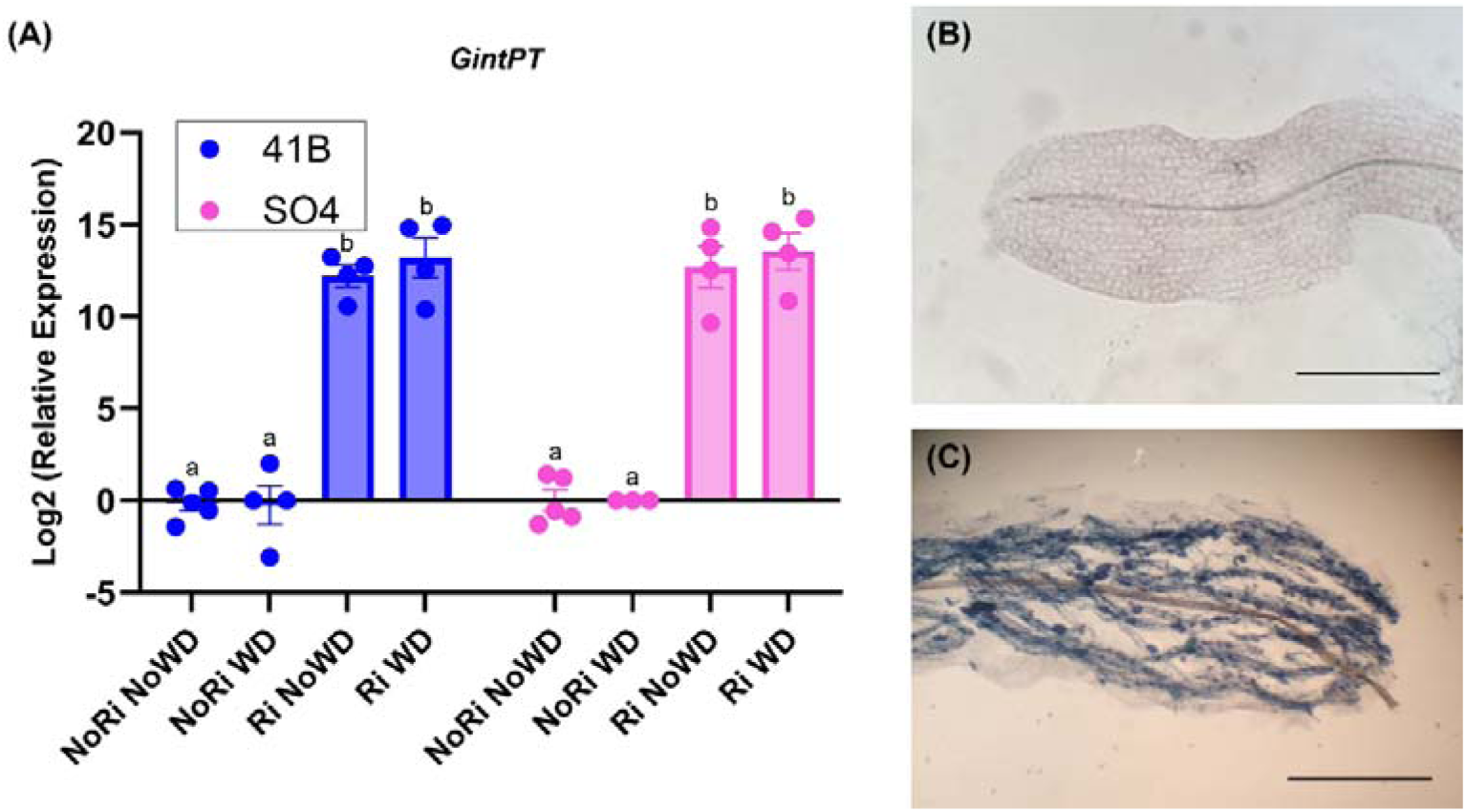
**Establishment and functionality of *R. irregularis* symbiosis in 41B and SO4 rootstocks under WD and NoWD regimes.** (A) Barplot shows the relative expression (log_2_) of *GintPT* for each condition in both genotypes; (B-C) Pictures represent root colonization under NoRi (B) and Ri (C) inoculation treatment. Pictures were taken using a Nikon ALPHAPHOT-2 microscope (x 100 magnification; bars = 600 µm), blue structures are AMF hyphae, vesicles and arbuscules colored with Sheaffer skrit ink.

**Table 1.** Mycorrhization parameters of grapevine rootstocks under NoWD and WD regimes. Based on Trouvelot method (1986), parameters include: F% (frequency of mycorrhizal colonization), M% (intensity of colonization within the root system), and A% (arbuscule abundance in the colonized root fragments). Values are mean *±* SD. Letters indicate significant differences between treatments according to Tukey’s HSD pairwise comparison (*p* < 0.05); *NC* = no comparison.

| Mycorrhizal colonization |  |  |  |
| --- | --- | --- | --- |
| *R. irregularis* inoculated plants |  |  |  |
|  | NoWD | WD | p |
| SO4 | *N* = 6 | *N* = 10 | ANOVA |
| F (%) | 100.00 ± 0.00 | 95.75 ± 10.21 | NC |
| M (%) | 85.67 ± 5.00<sup>a</sup> | 67.83 ± 16.22<sup>b</sup> | 2.15e<sup>-2</sup> |
| A (%) | 47.92 ± 19.92<sup>a</sup> | 20.74 ± 8.17<sup>b</sup> | 1.67e<sup>-3</sup> |
| 41B | *N* = 8 | *N* = 7 | ANOVA |
| F (%) | 100.00 ± 0.00 | 98.81 ± 3.15 | NC |
| M (%) | 74.52 ± 12.29<sup>a</sup> | 72.13 ± 7.71<sup>a</sup> | 6.67e<sup>-1</sup> |
| A (%) | 34.89 ± 15.73<sup>a</sup> | 38.75 ± 10.15<sup>a</sup> | 5.89e<sup>-1</sup> |

### 2. Symbiosis with *R. irregularis* enhanced growth parameters in both rootstocks, especially under water deficit

Growth was strongly affected by drought stress in both grapevine rootstock genotypes (Tab. 2). Indeed, in NoRi condition, drought-stressed plants showed a severe reduction in shoot elongation. With a mean height increase by only 3 cm for 41B and 4 cm for SO4, shoot elongation was reduced by 91.5% and 73% respectively. Symbiosis with *R. irregularis* significantly improved growth of both genotypes under drought conditions, reaching levels comparable to well-watered NoRi plants (Tab. 2). In 41B, *R. irregularis* inoculation also enhanced shoot growth in well-watered condition, since Ri–NoWD plants showed the highest elongation (42 cm), significantly higher than the control condition (NoRi–NoWD; 15 cm, p = 1.64^e-6^).

**Table 2.** Growth response of 41B and SO4 rootstocks colonized or not with *R. irregularis* under NoWD and WD regimes. Growth corresponds to the increase in shoot height measured as the difference between plant height at harvest and at the onset of drought stress. Values are mean *±* SD. Letters indicate significant differences between condition/treatment according to adequate post-hoc test: (p < 0,05).

| Plant Growth |  |  |  |  |  |
| --- | --- | --- | --- | --- | --- |
|  | NoRi |  | Ri |  | p |
|  | NoWD | WD | NoWD | WD |  |
| SO4 | $N = 10$ | $N = 10$ | $N = 6$ | $N = 10$ | Kruskal-Wallis |
| Growth (cm)<sup>(1)</sup> | $35 \pm 11^a$ | $3 \pm 1^b$ | $36 \pm 9^a$ | $24 \pm 11^a$ | 2.23E-05 |
| 41B | $N = 9$ | $N = 9$ | $N = 8$ | $N = 7$ | ANOVA |
| Growth (cm)<sup>(1)</sup> | $15 \pm 8^{bc}$ | $4 \pm 2^c$ | $42 \pm 10^a$ | $17 \pm 12^b$ | 5,12E-09 |

Dry biomass measurements revealed distinct responses of 41B and SO4 to water deficit and mycorrhization (Fig. 2). Whereas drought stress had no significant impact on aerial and root biomasses in 41B, it led to reduction of both aerial and root biomasses in SO4 (Fig. 2). In 41B, symbiosis with *R. irregularis* had a positive effect on aerial biomass, especially in well-watered condition. In SO4, symbiosis with Ri had more significant impact on both aerial and root biomasses, allowing to mitigate the effect of water deficit. Indeed, in SO4, leaf biomass of Ri-WD condition were similar to NoRi-NoWD (control) condition (Fig. 2). In addition, root biomass of Ri-WD condition was significantly higher compared to NoRi-WD (Fig.2). Mycorrhization also positively impacts aerial biomass of well-watered SO4 plants. Overall, these results indicate that *R. irregularis* symbiosis tended to confer beneficial effect on aerial biomass in both rootstock genotypes and conditions. In addition, a favorable effect of mycorrhization was evidenced on root biomass for SO4 only.

**Figure 2.**
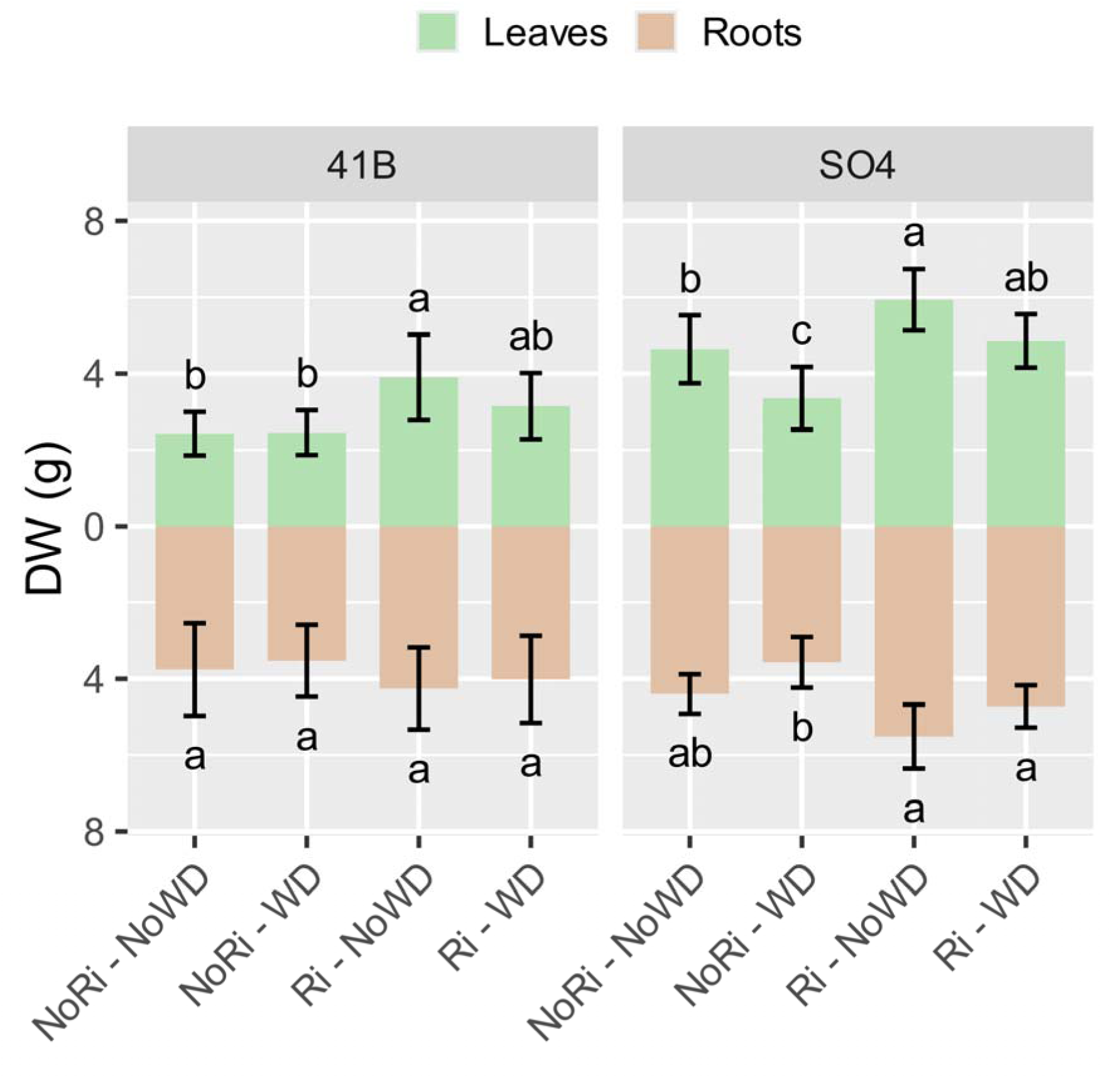
**Effect of *R. irregularis* symbiosis on dry weight (DW) of aerial and root parts in 41B and SO4 rootstocks under NoWD and WD regimes.** Upper panel shows aerial part DW biomass (light green) and lower panel root DW biomass (light brown). Each bar represents the mean ± SD, number of replicates is the same as for growth measurements. Letters indicate significant differences among conditions according to one-way ANOVA followed by Tukey’s HSD for 41B roots and SO4 leaves and Kruskal-Wallis followed by Dunn comparisons for 41B leaves and SO4 roots (p < 0,05).

### 3. Symbiosis with *R. irregularis* improved photosynthetic parameters under water deficit

Gas exchange and chlorophyll fluorescence measurements revealed strong effects of drought stress and *R. irregularis* symbiosis on the photosynthetic performance of both grapevine rootstock genotypes (Fig. 3). Drought stress significantly reduced several photosynthesis parameters in non-mycorrhized 41B and SO4: carbon net assimilation (A) (Fig. 3A), quantum yield of PSII (ϕPSII, Fig.3D) and stomatal conductance (gsw, Fig 3C) were significantly lower in NoRi-WD plants compared to NoRi-NoWD controls. However, the difference was not statistically significant for gsw in 41B, indicating a lowest stomatal closure in this genotype. Maximum quantum efficiency (Fv/Fm, Fig. 3B) followed the same trend, although not statistically significant. In water deficit condition and for both genotypes, mycorrhized plants (Ri-WD condition) maintained A, gsw, ΦPSII and Fv/Fm values comparable to those of well-watered plants (Ri-NoWD, p > 0.05). This suggests a positive effect of mycorrhization on water status and photosynthetic apparatus under drought conditions. In SO4, gsw in Ri-WD plants was slightly lower compared to Ri-NoWD, but the difference was not significant. Overall, these results indicate that drought stress negatively affected gas exchanges and photochemical efficiency, but *R. irregularis* symbiosis mitigated these effects in both 41B and SO4 rootstocks.

**Figure 3.**
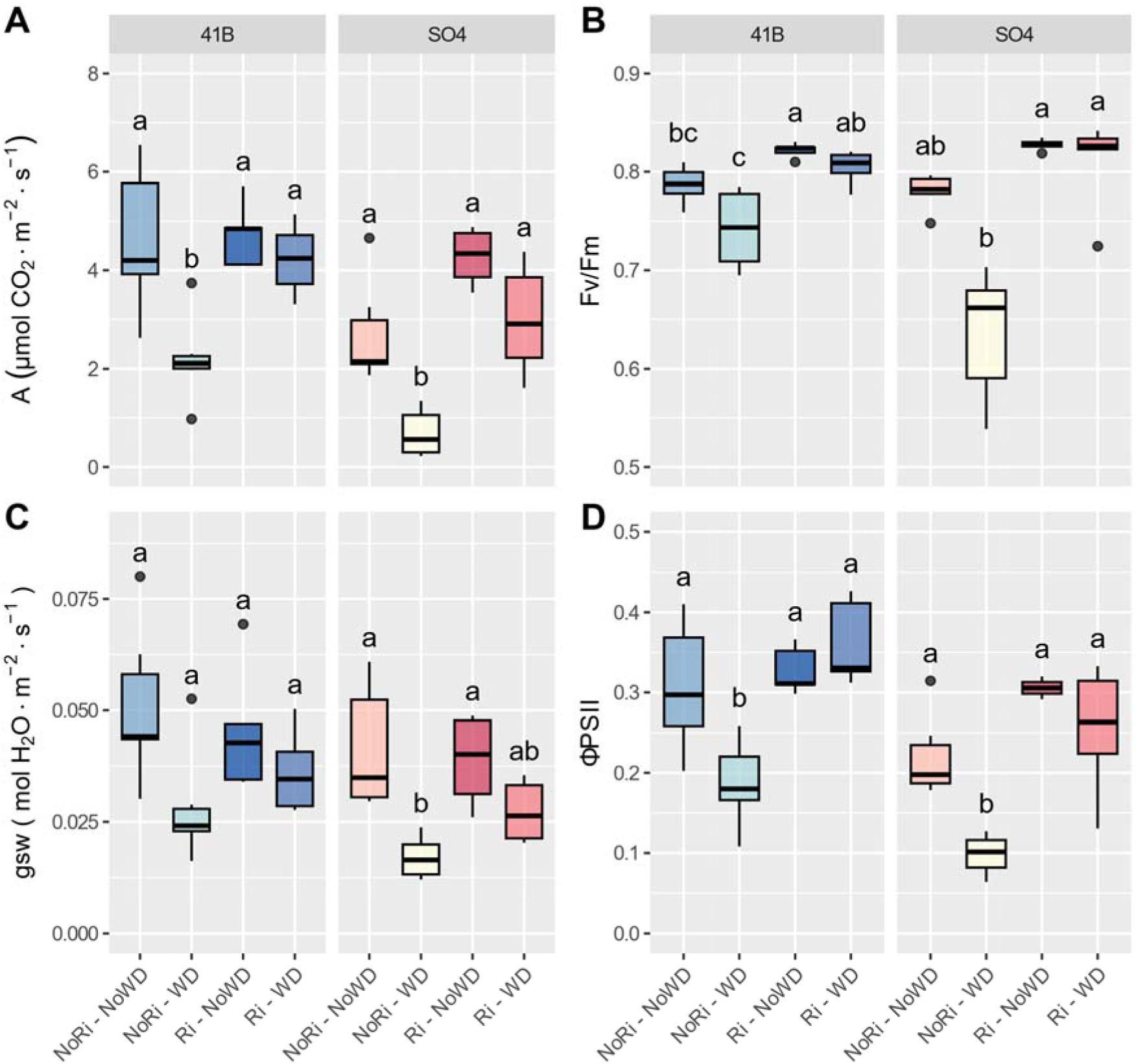
**Effect of *R. irregularis* symbiosis on gas exchanges and chlorophyll fluorescence in 41B and SO4 rootstocks under WD and NoWD regimes.** All parameters were measured by LI-6800 system (Li-COR®) on n = 4 – 6 biological replicates. (A) Carbon net assimilation; (B) Maximum quantum efficiency of photosystem II; (C) Stomatal conductance to water vapor ; (D) Quantum yield of photosystem II. Data are mean ± SD. Letters indicate significant differences between condition/treatment according to one-way ANOVA followed by Tukey’s HSD or Kruskal-Wallis followed by Dunn comparisons (p < 0,05).

### 4. Symbiosis with *R. irregularis* differentially impacts mineral elements in 41B and SO4, but enhances phosphorus levels in both rootstocks

Mineral element analysis revealed clear treatment-dependent variations in nutrient accumulation in leaves, roots and at the whole plant scale for both SO4 and 41B rootstock (Fig. 4). In general, more significant up-regulations (compared to control) of mineral elements were highlighted in SO4 and more downregulations appeared in 41B in the different conditions (Fig. 4). In both genotypes, AMF inoculation significantly increased phosphorus concentrations in roots and leaves, both in well-watered and water-deficit conditions (Tab. S3, S4). At the whole plant scale in SO4, P content in Ri plants increased by 57.7 % in NoWD and by 40.3 % in WD condition. Results were comparable in 41B where P increased by 57 % in Ri plants for NoWD condition and by 34 % in WD condition.

**Figure 4.**
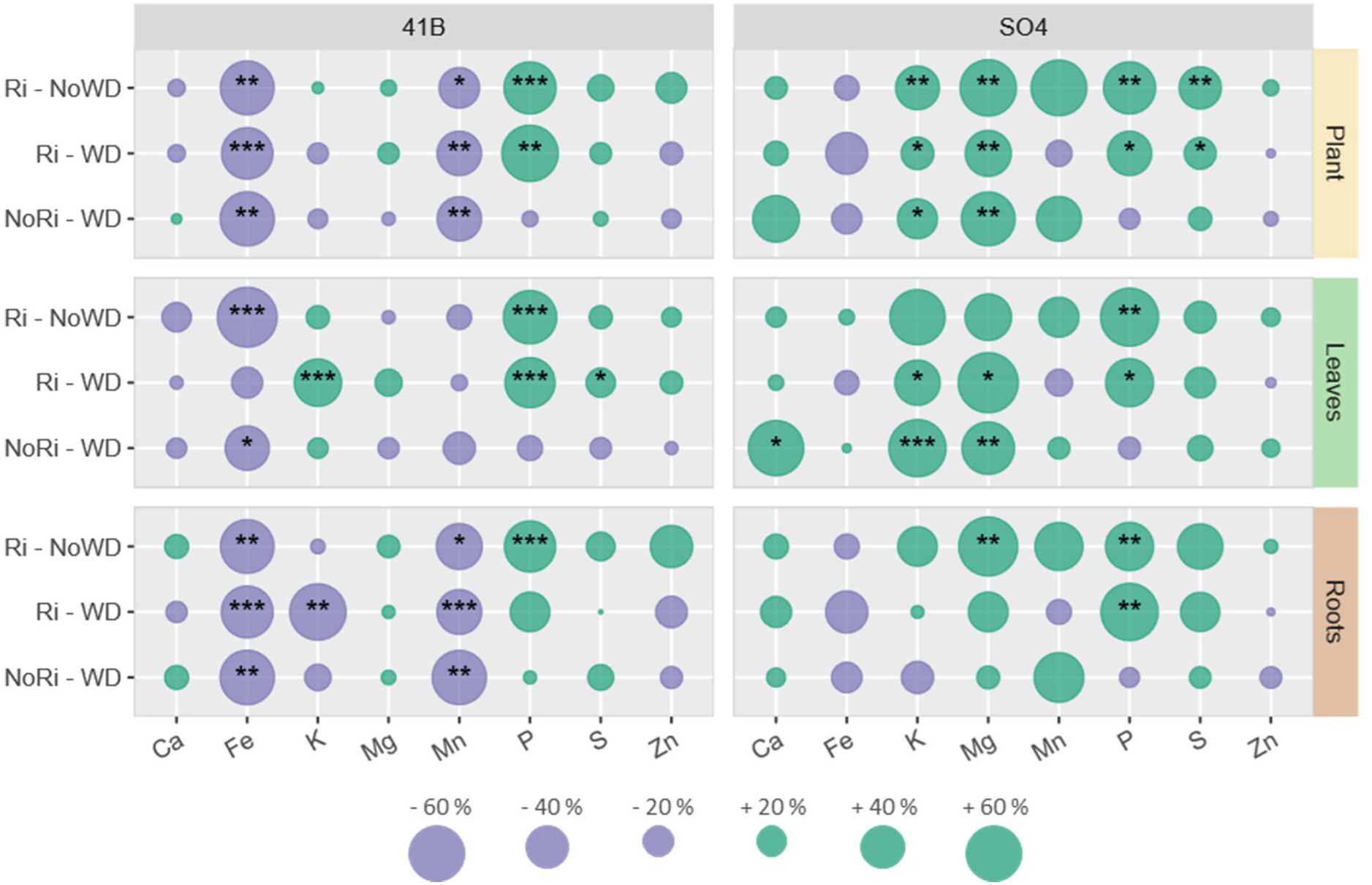
Effect of *R. irregularis* symbiosis on the elemental composition of roots, leaves and whole plant in 41B and SO4 rootstocks under NoWD and WD regimes. Bubble size represents the variation (%) of mineral content for each treatment/condition compared to content in NoRi-NoWD control. Bubble color represents the trend of element content variation compared to control: light green for positive difference (increase relative to NoRi-NoWD) and purple for negative differences (decrease relative to NoRi-NoWD). Asterisks show significant element content differences compared to control according to chosen pairwise comparison (* p-val < 0,05 ; ** p-val < 0,01 ; *** p-val < 0,001).

Concerning the other elements such as Fe, Mn, K, and Mg, both mycorrhization and water deficit differently affected their contents in the two genotypes (Fig.4). In 41B, Fe and Mn contents decreased significantly compared to the control condition as a result of mycorrhization and water limitation. These decreases concerned roots and leaves for Fe, but only roots for Mn. Concerning K, the content tended to respectively decrease in roots and increase in leaves following mycorrhization in water deficit condition in 41B.

In SO4, Fe contents also tended to decrease due to both mycorrhization and water deficit, although this effect only concerned roots and was not statistically significant. Water shortage significantly increased K content in leaves of both Ri and NoRi plants. In addition, the results show a positive effect of mycorrhization alone on K contents at the whole plant level (Fig.4). Mg content in whole SO4 plants significantly increased for the three conditions compared to NoRi-NoWD control, but differentially regarding the organ concerned. In roots, mycorrhization alone led to increase in Mg (37.7 %). Mg content also increased in leaves of SO4 as a result of water deficit. Finally, Ri-colonized SO4 whole plants also showed enhanced sulfur both in WD and NoWD conditions.

### 5. Symbiosis with *Rhizophagus irregularis* and water deficit differentially affect aquaporin gene expression in 41B and SO4 roots

Since aquaporins are good candidates to contribute to water uptake and drought tolerance in roots, we focused on the expression of genes encoding grapevine aquaporins representative from different families in roots of 41B and SO4. We studied the expression of four Plasma membrane Intrinsic Proteins (*VvPIP1.2*, *VvPIP1.4*, *VvPIP 2.1* and *VvPIP2.2*), two Tonoplast Intrinsic Proteins (*VvTIP1.1* and *VvTIP2.2*), two Nodulin26-like Intrinsic Proteins (*VvNIP1.1* and *VvNIP5.1*) and one uncategorized (X) Intrinsic Protein (*VvXIP*).

Contrasted results were obtained concerning aquaporin gene expression in 41B and SO4 roots under different water regime. Generally, expression of a higher number of aquaporin genes was up-regulated by mycorrhization in 41B compared to SO4 (Fig. S1). Expression of aquaporin genes significantly affected by mycorrhization and/or water deficit is detailed in Figure 5.

**Figure 5.**
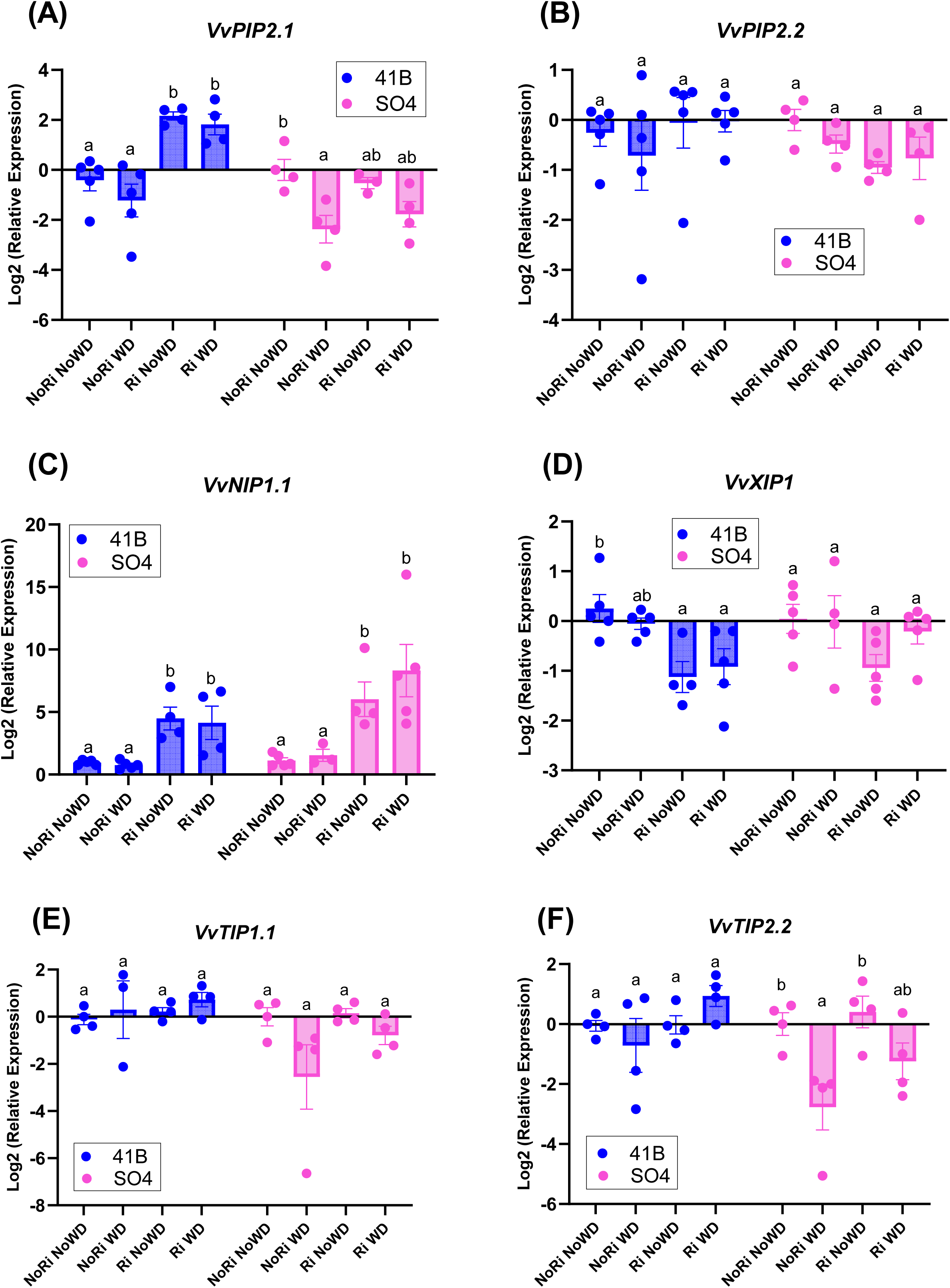
**Effect of *R. irregularis* symbiosis on the expression of root aquaporins in 41B and SO4 rootstocks under WD and NoWD regimes.** Barplots shows the relative expression (Log_2_) of Aquaporin genes in roots of both genotypes for each treatment/condition. Transcript levels were normalized to *V. vinifera ACTIN* and *EF1α* transcript levels. For each gene, relative expression was obtained with the 2^-ΔΔCT^ method and indicates mean normalized expression in the different conditions compared with normalized expression in the NoRi-NoWD condition (control). (A) *VvPIP2.1*, (B) *VvPIP2.2*, (C)*VvNIP1.1*, (D) *VvXIP1*, (E) *VvTIP1.1*, and (F) *VvTIP2.2*. Values are means ± SD of n = 3–5 biological replicates. Barplots were created using GraphPad Prism.

In 41B, water stress did not result in extensive impact on aquaporin expression. In contrast, Ri colonization significantly up-regulated the expression of *VvPIP2.1* and *VvNIP1.1*, both in well-watered and water stress conditions (Fig. 5A-C). In contrast, the expression of *VvXIP* was downregulated by Ri colonization in both conditions compared to NoRi-NoWD control (Fig. 5D). Tonoplast aquaporin (*VvTIP1.1* and *VvTIP2.2*) expression was not significantly affected in 41B in the different conditions.

In SO4, water stress triggered downregulation of the expression of *VvPIP2.1* in NoRi, and to a lesser extent, in Ri conditions (Fig. 5A). The expression of *VvPIP2.2* was also slightly lower (p=0.0996) in roots of Ri plants in well-watered condition (Figure 5B). As for 41B, mycorrhization up-regulated the expression of *VvNIP1.1* in both well-watered and water deficit conditions (Fig. 5C). In contrast to 41B, in SO4, *VvTIP1.1* and *VvTIP2.2* expression tended to be lower following water stress in non-mycorrhized plants (although only significantly for *VvTIP2.2*) (Fig. 5E-F). Finally, Ri colonization did not result in significant changes concerning *VvXIP* expression, although a tendency to downregulation was observed in Ri NoWD roots (Fig.5B).

## Discussion

### 1 Mycorrhization is differently affected by water stress in 41B and SO4 and could help the plant to sustain water limiting conditions

In this work, we explored the benefit of symbiosis with the AMF *Rhizophagus irregularis* on the tolerance of two widely-used rootstocks to moderate to severe water deficit. Few studies focused on rootstock behavior following colonization with a single AMF strain and in water shortage conditions. Since we used plants grown in inorganic substrate and watered with a nutrient solution, we are aware that water deficit also led to a reduced supply of mineral elements. However, this trend reflects natural conditions, where water deficit also results in less mineral element mobility and uptake by the plant (Ahanger et al. 2016).

Our results show that *Rhizophagus irregularis* established a successful symbiosis with both 41B and SO4 genotypes. However, under stressed condition, while water deficit had no effect on the frequency, intensity and functionality of the symbiosis in 41B, it significantly reduced both mycorrhizal intensity and arbuscule abundance in SO4. This result shows that the interaction between grapevine genotype and water stress impact symbiosis with *R. irregularis*. Although a higher arbuscular colonization has been described after the onset of deficit irrigation in grapevine (Schreiner et al. 2007), other studies also reported that water deficit resulted in decline in AMF colonization (Valentine et al. 2006; Trouvelot et al. 2015). However, we could notice that the percentage of arbuscules (20%) in SO4 roots submitted to water stress is significant. Despite reduction of M and A, we also observed a high expression of GintPT, which is a marker of symbiosis functionality (Duret et al. 2022), both in WD and NoWD in SO4, similar to what is observed in 41B. This result indicates that an active symbiosis occurs in the two rootstocks and regardless of the plant’s water status.

Concerning growth parameters, water deficit resulted in a strong inhibition of shoot elongation in both 41B and SO4, confirming the effect of drought. In contrast, its impact on leaf dry weight was less pronounced for the two genotypes and particularly for 41B. This result could be explained by the fact that a substantial part of leaf growth occurred before the onset of water stress. It also suggests that in 41B, although internode length is reduced, the numbers of internodes and leaves as well as leaf size are not significantly affected. Water deficit also led to decrease in root dry weight, but only in SO4. These results underline that drought more severely impacts growth of SO4 compared to 41B, in accordance with data reporting that SO4 is more sensitive to water depletion compared to 41B (Lovisolo et al. 2016).

Interestingly, under water deficit condition, symbiosis with *R. irregularis* allowed to restore all growth parameters (i.e., shoot elongation, leaf and root dry weights) to the levels observed in non-stressed plants for both genotypes. In the non-stressed condition, Ri had also a significant positive effect on shoot elongation in 41B and on leaf dry weights in both 41B and SO4. Overall, these results underline that although mycorrhization parameters seemed to be affected by water deficit in SO4, a positive effect of mycorrhization was noticed on the growth of both rootstocks, especially under stress condition, confirming the functionality of the symbiosis. These results on two widely used rootstocks are consistent with other studies reporting that mycorrhization could help grapevine to face the deleterious effect of drought, both in controlled and vineyard conditions. Indeed, Ye et al. (2022) showed that AMF colonization improves growth and relative water content of greenhouse grown *V. vinifera* cv. Ecolly under drought stress. Torres et al. (2021) further reported that AMF inoculation in vineyard improves 2-year-old Merlot grapevine vegetative growth, water status and photosynthetic activity in water limited conditions of a hyper arid growing season. Overall, this beneficial effect in water deficit condition could be explained by a better soil exploration and thus access to water in mycorrhized roots as well as a better nutrition and photosynthesis (see below).

In line with the results on growth, we observed a significant effect of water deficit on several photosynthesis parameters. In non-mycorrhized plants, net assimilation, maximum quantum efficiency (Fv/Fm) and quantum yield of photosystem II were lower in the WD condition compared to well-watered plants. It is known that water deficit impacts metabolism of chlorophyll pigments and limits water availability for photochemistry reactions (Flexas et al. 1999; Medrano et al. 2003). Fv/Fm (about 0.8 in healthy non-stressed plants) is a parameter sensitive to stress that affects PSII reaction centers. Concerning stomatal conductance, water deficit also resulted in a decrease in gsw in both genotypes (although only significant in SO4), underlying that rootstock response to drought involved stomatal closure to limit water loss. Similar physiological responses have been described in other grapevine cultivars such as Shiraz and Grenache under drought and heat stress (Cogato et al. 2022). In WD condition, symbiosis with *R. irregularis* improved all analyzed photosynthesis parameters, restoring them to levels similar to non-stressed plants. Mycorrhization had also a beneficial effect on maximum quantum efficiency in non-stressed 41B plants. Improvement of photosynthesis parameters in mycorrhized plants could be explained by a better access to water and nutrients that are essential to photosynthesis. Indeed, the improvement of water status in Ri-colonized plants could positively influence gas exchanges and the efficiency of photochemistry of PSII. Several studies reported that AMF symbiosis improve photosynthesis capacity and chlorophyll content in grapevine. Nikolaou et al. (2003) reported that *Funneliformis mosseae* improved leaf water potential and CO_2_ assimilation in Cabernet Sauvignon grafted on different rootstocks and submitted to drought. A positive effect of native vineyard AMF on photosynthesis was also outlined in Paulsen 1103 (Meyer 2020) and of the AMF *Glomus iranicum* in Crimson seedless (Nicolás et al. 2015). In addition, a recent study by Ye et al. (2022) reported enhanced photosynthetic activity and increased chlorophyll content in water-stressed *Vitis vinifera* cv. Ecolly following mycorrhization with a commercial inoculum containing four different AMF species.

### 2 *R. irregularis* beneficial effect could result from better grapevine mineral nutrition in stressed condition, with a contrasted effect in 41B and SO4

It is well known that mineral elements play central role in several physiological and biochemical processes through enzyme activation, osmoregulation, protein synthesis and photosynthesis.

Overall, the concentrations of the different nutrients in the rootstock leaves were within the same range as those reported in previous studies of greenhouse-grown grapevine cuttings cultivated in low-P substrates and inoculated with AMF (Moukarzel et al. 2022).

Generally, 41B and SO4 showed contrasting responses in terms of elemental contents to both water depletion and mycorrhization. Focusing on water depletion, drought significantly decreased Fe (roots and leaves) and Mn (roots) levels in non-mycorrhized 41B, whereas no significant effect was observed for the other elements. The decrease of mineral contents in roots may be a direct consequence of hydromineral deficiency associated with water depletion. Interestingly, this decrease is less pronounced in Ri plants. In SO4, water deficit did not lead to decrease in Fe and Mn. In contrast, in SO4, water stress resulted in increased levels of K, Ca and Mg in the leaves. Increased mineral contents could reflect a higher response to water depletion in this rootstock. Indeed, Mg is an important component of photosynthetic reactions through enzyme activation and is also involved in chlorophyll formation (Tian et al. 2021). The calcium ion Ca^2+^ also plays a central role in abiotic stress responses, by mediating hormonal responses leading to expression of drought responsive genes (Ahanger et al. 2016).

Concerning the effect of symbiosis with *R. irregularis*, a strong and consistent response was observed for phosphorus (P) in both rootstocks. Indeed, mycorrhized plants, and especially those in water-depleted condition, exhibited a clear increase in P contents, both in roots and leaves. P is one of the major macronutrients essential for plant development and metabolism. The significant effect of AMF symbiosis on P content is in accordance with literature and it is well-established that mycorrhiza improves P nutrition in plants. In the context of water limitation, plants colonized by AMF can also obtain phosphorus through the mycorrhizal pathway, which is frequently more efficient in mobilizing soil phosphorus than direct root uptake (Smith et al. 2011; Püschel et al. 2021). In grapevine, an increase in P levels in roots and/or leaves was evidenced in almost all studies of grapevine inoculated with AMF (Schreiner 2005). Less data are available on the effect of mycorrhiza on grapevine nutrition in drought conditions, however, Cardinale et al. (2022) reported that mycorrhized 1103-Paulsen rooted cuttings accumulated higher leaf mineral element (including P) under exacerbated summer conditions. Nikolaou et al. (2003) also found higher P contents in leaves of Cabernet-Sauvignon grafted on 41B and exposed to drought. In this study, enhanced P contents in mycorrhized roots are in accordance with high level of *GintPT* expression, which could be involved in both P acquisition from soil and transfer towards plant cells (Fiorilli et al. 2013). The beneficial outcome of Ri symbiosis on P is likely responsible for photosynthesis improvement in water deficit condition, since it is well established that adequate supply of P is crucial for photosynthetic carbon assimilation. Indeed, P is necessary for co-factors (ATP, NADPH) and phosphorylated intermediate production in the Calvin cycle. An inhibition of photosynthesis triggered by inadequate Pi nutrition has been reported in a number of plant species (Pessarakli 1996).

The effect of mycorrhization on other mineral elements seems to be more genotype-dependent. Ri colonization tended to decrease levels of metal elements such as Fe and Mn, especially in the roots and leaves of 41B. This result contrasts with other studies showing increase of Mn after colonization with AMF (Moukarzel et al. 2022). However, a negative effect of mycorrhization on root metal contents, especially Fe, has been reported in several studies (Biricolti et al. 1997). Since metals exhibit toxicity towards microorganisms, it is known that metals could be detoxified and transformed by fungi, including mycorrhizas (Gadd 2010). Furthermore, the observed improved uptake of certain nutrients in roots of mycorrhized plants may lead to competitive effects that reduce the uptake or translocation of other elements, modifying the balance of ion uptake (Ferrol et al. 2016).

Contrasted response to AMF colonization in the two genotypes was also evidenced concerning K, Ca, Mg and S levels. In SO4, water deficit and mycorrhization led to higher K in the leaves and the whole plant respectively. In 41B water deficit condition, mycorrhization resulted in lower K contents in roots, whereas leaf levels were increased. Potassium is one of the most abundant elements involved in a number of physiological processes, especially photosynthesis through regulation of RuBiSO activity and regulation of stomatal movement. In addition, K is involved in response to abiotic stresses thanks to osmoregulation and ROS detoxification in the leaves (Ahanger et al. 2016; Rawat et al. 2022). In mycorrhized 41B, K may be remobilized from roots to shoots under water stress, thereby contributing to the mitigation of drought stress. AMF colonization had also a positive effect on Mg levels in both roots and leaves, as well as on S content at the whole plant level in SO4 but not in 41B. As component of the chlorophyll molecule and co-factor of many enzymes, Mg is an important component involved in photosynthesis regulation and protein synthesis (Ahanger et al. 2016). Sulfur is essential as part of protein disulfide bonds, ammino acids and co-factors and sulfur-containing molecules are involved in stress signaling (Narayan et al. 2023). Many reports show that plant AMF colonization improves sulfur content by increasing its uptake from the soil through modulation of plant sulfate transporter’s expression (Narayan et al. 2023).

In conclusion, AMF symbiosis seems to result in more element up-regulations in SO4 compared to 41B, in accordance with significant positive effect on growth in this genotype in stress condition, underlying the interest of mycorrhization for drought sensitive rootstocks.

### 3 Symbiosis with *R. irregularis* resulted in contrasted regulation of aquaporin expression in roots of 41B and SO4

Aquaporins (AQP) are small channels involved in the passive movement of water and small molecules across membranes (Maurel et al. 2015). In the grapevine genome, Shelden et al. (2009) identified 23 genes encoding Major Intrinsic Proteins (MIPs) belonging to PIP (Plasma Membrane intrinsic proteins, split in two subgroups VvPIP1s and VvPIP2s), TIP (Tonoplast Intrinsic proteins), NIPs (nodulin26-like Intrinsic Proteins), SIPs (small basic intrinsic proteins) and XIPs (uncategorized intrinsic proteins) (Maurel et al. 2015). AQPs play a central role in water status regulation in grapevine since their modulation has been involved in cell-to-cell water radial transport from soil to root xylem vessels, regulation of cell hydraulic conductance and water status maintenance (Sabir et al. 2021; Braidotti et al. 2024). During drought stress, AQP expression from PIP and TIP families has been involved in the regulation of hydraulic conductivity and thus appears as interesting candidates for grapevine adaptation to water deficit (Gambetta et al. 2012, 2017; Vandeleur et al. 2014; Labarga et al. 2023). Mycorrhizal symbiosis could also influence cell-to-cell water and solute flux through aquaporins in plants (Ruiz-Lozano and Aroca 2017). However, in grapevine, data concerning aquaporin expression following AMF colonization combined with drought conditions are scarce. In a recent study, Guan et al. (2025) revealed that aquaporin expression is strongly associated with recovering after water stress in cv. Pinot Noir, highlighting their importance as regulator of water homeostasis in grapevine. Our data revealed that in both rootstock genotypes, mycorrhization led to significant higher expression of *VvNIP1.1* both in well-watered and drought condition. NIP1.1 is a member of nodulin26-like Intrinsic Protein showing significant water transport activity (Sabir et al. 2021) and enhanced *VvNIP1.1* expression has been reported in several mycorrhized rootstocks (Sportes et al. 2023). Higher *VvNIP1.1* expression evidenced in mycorrhized roots under water deficit suggests that symbiosis improves root water transport in both rootstocks. In contrast, *VvXIP* expression is negatively regulated by mycorrhization in both WD and NoWD conditions, but only significantly in 41B. VvXIP1, localized within the endoplasmic reticulum membrane, is unable to transport water (Noronha et al. 2016). This aquaporin is permeable to glycerol, hydrogen peroxide, heavy metals and metalloids. Interestingly, Noronha et al. (2016) also reported downregulation of *VvXIP1* in leaves of potted vines under water deficit , suggesting its possible involvement in osmotic regulation. In our study, downregulation of *VvXIP1* in mycorrhized 41B roots also suggests a link with downregulation of metal transport.

Mycorrhization also triggered higher expression of *VvPIP2.1* in 41B roots, both in control and drought conditions. In contrast, in SO4, *VvPIP2.1* expression did not change significantly following Ri colonization, but was repressed by water deficit, both in non-mycorrhized and mycorrhized roots. *VvPIP2.1* has been characterized as a highly water permeable aquaporin (Shelden et al. 2009). Interestingly, constitutive higher expression of *VvPIP2.1* and *VvPIP2.2* were evidenced in rootstock genotypes (110R and 140R) known as more tolerant to water deficit (Gambetta et al. 2012). In addition, Bonarota et al. (2024) revealed that higher performance to water deficit of a drought-tolerant rootstock (*Vitis champigni*) involved higher expression of *VvPIP2.1*. In our study, expression of another *PIP*, *VvPIP2.2,* was not significantly impacted by AMF symbiosis and/or water deficit in 41B, whereas it was slightly downregulated by water deficit in SO4. The same trend was observed for *TIP* expression: whereas water deficit had no significant impact on both *VvTIP1.1* and *VvTIP2.2* expression in 41B, it had a negative impact, especially on *VvTIP2.2* expression in mycorrhized and non-mycorrhized SO4 roots. TIP are tonoplast water permeable AQP that could allow cells to adjust osmotic fluctuations triggered by water deficit in the cytosol (Kjellbom et al. 1999). Downregulation of *VvPIP2.1* and *VvTIP2.2* was observed in water-stressed roots of Touriga National, in line with decrease of root hydraulic conductivity (Zarrouk et al. 2016). Galaz et al. (2024) also recently underlined that *VvTIP1.1* and *VvTIP2.1* expression is correlated to stomatal conductance response in Cabernet Sauvignon grafted on different rootstocks and subjected to water scarcity.

Overall, lower expression of several AQPs in SO4 could result from higher impact of drought stress on SO4 root cells. Indeed, lower root biomass resulting from drought has only been shown in this rootstock. Conversely, constitutive overexpression of *VvPIP2.1* following mycorrhization in 41B could confer better water uptake and tolerance to water deficit.

## Conclusions

This study highlights global positive effect of symbiosis with *Rhizophagus irregularis* on growth and photosynthesis parameters in two rootstocks widely used in France Northeast vineyards, with a more pronounced positive effect however in SO4. Our results further emphasize that interaction between AMF and rootstock genotype induces contrasted effects on grapevine nutrition. Whereas *R. irregularis* had higher positive effect on mineral nutrition in SO4, it had stronger positive effect on root aquaporin expression in 41B. It thus appears that nursery plant mycorrhization could be of interest for rootstocks sensitive to drought. However, attention should be paid to the effect of mycorrhizal inoculation on endemic AMF communities and future studies will address this issue.

## Acknowledgments

We are grateful to the Agronutrition company (Carbonne, France) for providing the *R. irregularis* inoculum. We thank Dr. Marc Lollier for help in statistical analysis. We are grateful to Lucas Charrois (Lorraine University, Soil and Environment Laboratory) for element quantification with ICP-OES.

## Author contributions

Conceived and designed experiments: JC, LDB, LV, LY

Performed experiments: LG, ML, LY, HL, LDB, LV, JC

Analyzed experiments: LG, LY, ML, LV, LDB, JC

Wrote the paper: JC, LG, LY, LV

Funding acquisition: JC, LY

## References

Aguilera P, Ortiz N, Becerra N, et al (2022) Application of arbuscular mycorrhizal fungi in vineyards: water and biotic stress under a climate change scenario: new challenge for Chilean grapevine crop. Front Microbiol 13:826571

Ahanger MA, Morad-Talab N, Abd-Allah EF, et al (2016) Plant growth under drought stress: Significance of mineral nutrients. In: Ahmad P (ed) Water Stress and Crop Plants, 1st edn. Wiley, pp 649–668

Bernardo S, Marguerit E, Ollat N, et al (2025) Root system ideotypes: what is the potential for breeding drought-tolerant grapevine rootstocks? J Exp Bot 76:2970–2984. 10.1093/jxb/eraf006

Biricolti S, Ferrini F, Rinaldelli E, et al (1997) VAM Fungi and Soil Lime Content Influence Rootstock Growth and Nutrient Content. Am J Enol Vitic 48:93–99. 10.5344/ajev.1997.48.1.93

Bonarota M-S, Toups HS, Bristow ST, et al (2024) Drought response and recovery mechanisms of grapevine rootstocks grafted to a common *Vitis vinifera* scion. Plant Stress 11:100346. 10.1016/j.stress.2024.100346

Braidotti R, Falchi R, Calderan A, et al (2024) Multi-hormonal analysis and aquaporins regulation reveal new insights on drought tolerance in grapevine. J Plant Physiol 296:154243. 10.1016/j.jplph.2024.154243

Cardinale M, Minervini F, De Angelis M, et al (2022) Vineyard establishment under exacerbated summer stress: effects of mycorrhization on rootstock agronomical parameters, leaf element composition and root-associated bacterial microbiota. Plant Soil 478:613–634. 10.1007/s11104-022-05495-1

Chaves MM, Zarrouk O, Francisco R, et al (2010) Grapevine under deficit irrigation: hints from physiological and molecular data. Ann Bot 105:661–676

Cogato A, Jewan SYY, Wu L, et al (2022) Water Stress Impacts on Grapevines (*Vitis vinifera* L.) in Hot Environments: Physiological and Spectral Responses. Agronomy 12:1819. 10.3390/agronomy12081819

Dagher D, Taskos D, Mourouzidou S, Monokrousos N (2025) Microbial-Enhanced Abiotic Stress Tolerance in Grapevines: Molecular Mechanisms and Synergistic Effects of Arbuscular Mycorrhizal Fungi, Plant Growth-Promoting Rhizobacteria, and Endophytes. Horticulturae 11:592

Degu A, Hochberg U, Wong DCJ, et al (2019) Swift metabolite changes and leaf shedding are milestones in the acclimation process of grapevine under prolonged water stress. BMC Plant Biol 19:69. 10.1186/s12870-019-1652-y

Dietz K-J, Zörb C, Geilfus C-M (2021) Drought and crop yield. Plant Biol 23:881–893. 10.1111/plb.13304

Duret M, Zhan X, Belval L, et al (2022) Use of a RT-qPCR Method to Estimate Mycorrhization Intensity and Symbiosis Vitality in Grapevine Plants Inoculated with *Rhizophagus irregularis*. Plants 11:3237. 10.3390/plants11233237

Ferrol N, Tamayo E, Vargas P (2016) The heavy metal paradox in arbuscular mycorrhizas: from mechanisms to biotechnological applications. J Exp Bot 67:6253–6265. 10.1093/jxb/erw403

Fiorilli V, Lanfranco L, Bonfante P (2013) The expression of GintPT, the phosphate transporter of *Rhizophagus irregularis*, depends on the symbiotic status and phosphate availability. Planta 237:1267–1277. 10.1007/s00425-013-1842-z

Flexas J, Escalona JM, Medrano H (1999) Water stress induces different levels of photosynthesis and electron transport rate regulation in grapevines. Plant Cell Environ 22:39–48. 10.1046/j.1365-3040.1999.00371.x

Gadd GM (2010) Metals, minerals and microbes: geomicrobiology and bioremediation. Microbiology 156:609–643. 10.1099/mic.0.037143-0

Galaz A, Pérez-Donoso AG, Gambardella M (2024) Leaf Aquaporin Expression in Grafted Plants and the Influence of Genotypes and Scion/Rootstock Combinations on Stomatal Behavior in Grapevines Under Water Deficit. Plants 13:3427. 10.3390/plants13233427

Gambetta GA, Herrera JC, Dayer S, et al (2020) The physiology of drought stress in grapevine: towards an integrative definition of drought tolerance. J Exp Bot 71:4658–4676

Gambetta GA, Knipfer T, Fricke W, McElrone AJ (2017) Aquaporins and Root Water Uptake. In: Chaumont F, Tyerman SD (eds) Plant Aquaporins: From Transport to Signaling. Springer International Publishing, Cham, pp 133–153

Gambetta GA, Manuck CM, Drucker ST, et al (2012) The relationship between root hydraulics and scion vigour across *Vitis* rootstocks: what role do root aquaporins play? J Exp Bot 63:6445–6455. 10.1093/jxb/ers312

Goddard M-L, Belval L, Martin IR, et al (2021) Arbuscular Mycorrhizal Symbiosis Triggers Major Changes in Primary Metabolism Together With Modification of Defense Responses and Signaling in Both Roots and Leaves of *Vitis vinifera*. Front Plant Sci 12:721614. 10.3389/fpls.2021.721614

Guan L, Valenzuela AV, Sharma G, et al (2025) Aquaporin translation tunes plant water transport to external conditions in grapevine. Plant Physiol Biochem 110298

Ju Y, Yue X, Zhao X, et al (2018) Physiological, micro-morphological and metabolomic analysis of grapevine (*Vitis vinifera* L.) leaf of plants under water stress. Plant Physiol Biochem 130:501–510. 10.1016/j.plaphy.2018.07.036

Kjellbom P, Larsson C, Johansson I, et al (1999) Aquaporins and water homeostasis in plants. Trends Plant Sci 4:308–314

Kokkoris V, Banchini C, Paré L, et al (2024) Rhizophagus irregularis, the model fungus in arbuscular mycorrhiza research, forms dimorphic spores. New Phytol 242:1771–1784. 10.1111/nph.19121

Kozikova D, Pascual I, Goicoechea N (2024) Arbuscular Mycorrhizal Fungi Improve the Performance of Tempranillo and Cabernet Sauvignon Facing Water Deficit under Current and Future Climatic Conditions. Plants 13:1155. 10.3390/plants13081155

Labarga D, Mairata A, Puelles M, et al (2023) The Rootstock Genotypes Determine Drought Tolerance by Regulating Aquaporin Expression at the Transcript Level and Phytohormone Balance. Plants 12:718. 10.3390/plants12040718

Leeuwen C van, Darriet P (2016) The Impact of Climate Change on Viticulture and Wine Quality. J Wine Econ 11:150–167. 10.1017/jwe.2015.21

Lovisolo C, Lavoie-Lamoureux A, Tramontini S, Ferrandino A (2016) Grapevine adaptations to water stress: new perspectives about soil/plant interactions. Theor Exp Plant Physiol 28:53–66. 10.1007/s40626-016-0057-7

Maurel C, Boursiac Y, Luu D-T, et al (2015) Aquaporins in Plants. Physiol Rev 95:1321–1358. 10.1152/physrev.00008.2015

Medrano H, Escalona JM, Cifre J, et al (2003) A ten-year study on the physiology of two Spanish grapevine cultivars under field conditions: effects of water availability from leaf photosynthesis to grape yield and quality. Funct Plant Biol 30:607–619. 10.1071/FP02110

Meyer E (2020) Growth, heavy metal uptake, and photosynthesis in “Paulsen 1103” (*Vitis berlandieri* x *rupestris*) grapevine rootstocks inoculated with arbuscular mycorrhizal fungi from vineyard soils with high copper contents. Vitis J Grapevine Res. 10.5073/VITIS.2020.59.169-180

Moukarzel R, Ridgway HJ, Liu J, et al (2022) AMF Community Diversity Promotes Grapevine Growth Parameters under High Black Foot Disease Pressure. J Fungi 8:250. 10.3390/jof8030250

Nakashima K, Yamaguchi-Shinozaki K (2013) ABA signaling in stress-response and seed development. Plant Cell Rep 32:959–970. 10.1007/s00299-013-1418-1

Narayan OP, Kumar P, Yadav B, et al (2023) Sulfur nutrition and its role in plant growth and development. Plant Signal Behav 18:2030082. 10.1080/15592324.2022.2030082

Nicolás E, Maestre-Valero JF, Alarcón JJ, et al (2015) Effectiveness and persistence of arbuscular mycorrhizal fungi on the physiology, nutrient uptake and yield of Crimson seedless grapevine. J Agric Sci 153:1084–1096. 10.1017/S002185961400080X

Nikolaou N, Angelopoulos K, Karagiannidis N (2003) Effects of drought stress on mycorrhizal and non-mycorrhizal cabernet sauvignon grapevine, grafted onto various rootstocks. Exp Agric 39:241–252. 10.1017/S001447970300125X

Nogales A, Rottier E, Campos C, et al (2021) The effects of field inoculation of arbuscular mycorrhizal fungi through rye donor plants on grapevine performance and soil properties. Agric Ecosyst Environ 313:107369. 10.1016/j.agee.2021.107369

Noronha H, Araújo D, Conde C, et al (2016) The Grapevine Uncharacterized Intrinsic Protein 1 (VvXIP1) Is Regulated by Drought Stress and Transports Glycerol, Hydrogen Peroxide, Heavy Metals but Not Water. PLOS ONE 11:e0160976. 10.1371/journal.pone.0160976

Pessarakli M (1996) Handbook of Photosynthesis, Second Edition. CRC Press

Püschel D, Bitterlich M, Rydlová J, Jansa J (2021) Drought accentuates the role of mycorrhiza in phosphorus uptake. Soil Biol Biochem 157:108243. 10.1016/j.soilbio.2021.108243

Rawat J, Pandey N, Saxena J (2022) Role of Potassium in Plant Photosynthesis, Transport, Growth and Yield. In: Iqbal N, Umar S (eds) Role of Potassium in Abiotic Stress. Springer Nature Singapore, Singapore, pp 1–14

Raza A, Razzaq A, Mehmood SS, et al (2019) Impact of Climate Change on Crops Adaptation and Strategies to Tackle Its Outcome: A Review. Plants 8:34. 10.3390/plants8020034

Ruiz-Lozano JM, Aroca R (2017) Plant Aquaporins and Mycorrhizae: Their Regulation and Involvement in Plant Physiology and Performance. In: Chaumont F, Tyerman SD (eds) Plant Aquaporins: From Transport to Signaling. Springer International Publishing, Cham, pp 333–353

Sabir F, Zarrouk O, Noronha H, et al (2021) Grapevine aquaporins: Diversity, cellular functions, and ecophysiological perspectives. Biochimie 188:61–76. 10.1016/j.biochi.2021.06.004

Schmittgen TD, Livak KJ (2008) Analyzing real-time PCR data by the comparative CT method. Nat Protoc 3:1101–1108. 10.1038/nprot.2008.73

Schreiner RP (2005) Mycorrhizas and mineral acquisition in grapevines. In: Proceedings of the soil environment and vine mineral nutrition symposium. American Society for Enology and Viticulture Davis, CA, pp 49–60

Schreiner RP, Tarara JM, Smithyman RP (2007) Deficit irrigation promotes arbuscular colonization of fine roots by mycorrhizal fungi in grapevines (*Vitis vinifera* L.) in an arid climate. Mycorrhiza 17:551–562. 10.1007/s00572-007-0128-3

Shelden MC, Howitt SM, Kaiser BN, Tyerman SD (2009) Identification and functional characterisation of aquaporins in the grapevine, *Vitis vinifera*. Funct Plant Biol 36:1065–1078

Shi J, Wang X, Wang E (2023) Mycorrhizal Symbiosis in Plant Growth and Stress Adaptation: From Genes to Ecosystems. Annu Rev Plant Biol 74:569–607. 10.1146/annurev-arplant-061722-090342

Smith SE, Jakobsen I, Grønlund M, Smith FA (2011) Roles of Arbuscular Mycorrhizas in Plant Phosphorus Nutrition: Interactions between Pathways of Phosphorus Uptake in Arbuscular Mycorrhizal Roots Have Important Implications for Understanding and Manipulating Plant Phosphorus Acquisition. Plant Physiol 156:1050–1057. 10.1104/pp.111.174581

Songy A, Fernandez O, Clément C, et al (2019) Grapevine trunk diseases under thermal and water stresses. Planta 249:1655–1679. 10.1007/s00425-019-03111-8

Sportes A, Hériché M, Mounier A, et al (2023) Comparative RNA sequencing-based transcriptome profiling of ten grapevine rootstocks: shared and specific sets of genes respond to mycorrhizal symbiosis. Mycorrhiza 33:369–385. 10.1007/s00572-023-01119-3

Tian X-Y, He D-D, Bai S, et al (2021) Physiological and molecular advances in magnesium nutrition of plants. Plant Soil 468:1–17. 10.1007/s11104-021-05139-w

Torres N, Antolín MC, Goicoechea N (2018) Arbuscular Mycorrhizal Symbiosis as a Promising Resource for Improving Berry Quality in Grapevines Under Changing Environments. Front Plant Sci 9:897. 10.3389/fpls.2018.00897

Torres N, Yu R, Kurtural SK (2021) Arbuscular mycrorrhizal fungi inoculation and applied water amounts modulate the response of young grapevines to mild water stress in a hyper-arid season. Front Plant Sci 11:622209

Tramontini S, Vitali M, Centioni L, et al (2013) Rootstock control of scion response to water stress in grapevine. Environ Exp Bot 93:20–26. 10.1016/j.envexpbot.2013.04.001

Trouvelot A, Kough JL, Gianinazzi-Pearson V (1986) Mesure du taux de mycorhization VA d’un système radiculaire. Recherche de méthode d’estimation ayant une signification fonctionnelle. pp 217–221

Trouvelot S, Bonneau L, Redecker D, et al (2015) Arbuscular mycorrhiza symbiosis in viticulture: a review. Agron Sustain Dev 35:1449–1467. 10.1007/s13593-015-0329-7

Valat L, Deglène-Benbrahim L, Kendel M, et al (2018) Transcriptional induction of two phosphate transporter 1 genes and enhanced root branching in grape plants inoculated with Funneliformis mosseae. Mycorrhiza 28:179–185. 10.1007/s00572-017-0809-5

Valentine AJ, Mortimer PE, Lintnaar M, Borgo R (2006) Drought responses of arbuscular mycorrhizal grapevines. Symbiosis 41:127–133

Vandeleur RK, Sullivan W, Athman A, et al (2014) Rapid shoot-to-root signalling regulates root hydraulic conductance via aquaporins. Plant Cell Environ 37:520–538. 10.1111/pce.12175

Velaz M, Santesteban LG, Torres N (2025) Mycorrhizae and grapevines: the known unknowns of their interaction for wine growers’ challenges. J Exp Bot 76:3001–3015. 10.1093/jxb/eraf081

Vierheilig H, Schweiger P, Brundrett M (2005) An overview of methods for the detection and observation of arbuscular mycorrhizal fungi in roots. Physiol Plant 125:393–404. 10.1111/j.1399-3054.2005.00564.x

Wolkovich EM, Cook BI, Cortázar-Atauri IG de, et al (2025) Uneven impacts of climate change around the world and across the annual cycle of winegrapes. PLOS Clim 4:e0000539. 10.1371/journal.pclm.0000539

Ye Q, Wang H, Li H (2023) Arbuscular Mycorrhizal Fungi Enhance Drought Stress Tolerance by Regulating Osmotic Balance, the Antioxidant System, and the Expression of Drought-Responsive Genes in *Vitis vinifera* L. Aust J Grape Wine Res 2023:7208341. 10.1155/2023/7208341

Ye Q, Wang H, Li H (2022) Arbuscular mycorrhizal fungi improve growth, photosynthetic activity, and chlorophyll fluorescence of *Vitis vinifera* L. cv. Ecolly under drought stress. Agronomy 12:1563

Yung L, Sirguey C, Azou-Barré A, Blaudez D (2021) Natural Fungal Endophytes From *Noccaea caerulescens* Mediate Neutral to Positive Effects on Plant Biomass, Mineral Nutrition and Zn Phytoextraction. Front Microbiol 12:689367. 10.3389/fmicb.2021.689367

Zarrouk O, Garcia-Tejero I, Pinto C, et al (2016) Aquaporins isoforms in cv. Touriga Nacional grapevine under water stress and recovery—Regulation of expression in leaves and roots. Agric Water Manag 164:167–175. 10.1016/j.agwat.2015.08.013

