## Supplementary Information for "Symbiosis with *Rhizophagus irregularis* improves grapevine rootstock performances under water deficit conditions"

| | | Genes | Accession number | Forward primer/ Reverse Primer (5’→3’) | Reference |
|---|---|---|---|---|---|
| Grapevine | Reference genes | *VvAct* | XM_002282480.2 | TGCTATCCTTCGTCTTGACCTTG/ GGACTTCTGGACAACGGAATCTC | (Reid et al. 2006) |
| | | *VvEF1α* | CB977561 | GGACCGGGTTCATGAGCTGTTGG/ TGAATGCAGACTCGCCAGCGGT | (Reid et al. 2006) |
| | Genes encoding aquaporins | *VvNIP5.1* | MN723561.1 | CGCAGCACAAGTTTCAGCAT/ GCGGTGACAACGAACAAGAG | |
| | | *VvNIP1.1* | NM723560.1 | CTTGCAGTCCTTTGTCATCG/ ACTCCTTGCTGGATTCA TCG | |
| | | *VvPIP2.1* | NM_001280993.1 | TCCTTCTCCGCCAAGGACTA/ AGACCAAGCAATGCCCAGAA | (Zarrouk et al. 2016) |
| | | *VvPIP2.2* | DQ834699 EC969993 | TCCGCCAAGGACTATCATGAC/ CGCAATCAGAGCCCTGTAGAA | |
| | | *VvPIP1.2* | EF364433 | GCCATCGTCTACAACAAAG/ CAGGCTCTGGTCTTGAATG | |
| | | *VvPIP1.4* | DQ834697.1 EF364435 | TCTGTTTCTTCTTTTATTTGCTGCT/ ATTCAAAAGCTGCCCATTGT | |
| | | *VvXIP* | F6I152 | ATCATGTCGGTTGTTGTTGC/ CAGCGCGTGAGAAAGAGAT | |
| | | *VvTIP1.1* | KJ697717 | AGCCTTTATTGGCGGACACA/ GTAAACCAGGCCGAAGGTCA | (Zarrouk et al. 2016) |
| | | *VvTIP2.2* | KJ697718 | CCATTGTTGCTTGCCTTCTCC/ TTGGCACCCACTATGAACCC | (Zarrouk et al. 2016) |
| *R. irregularis* | | *GintPT* | AY037894 | AACACGATGTCAACAAAGCAAC/ AAGACCGATTCCATAAAAGCA | (Fiorilli et al. 2013) |

**Table S1.** List of primer pairs used for RT-qPCR.

Whole plant mineral content

| Characteristic | | Ri | | NoRi | | |
|---|---|---|---|---|---|---|
| | | **NoWD** | **WD** | **NoWD** | **WD** | **P** |
| Ca | SO4 | $^{a}$31 313 ± 2 038 | $^{a}$31 821 ± 3 620 | $^{a}$28 907 ± 3 790 | $^{a}$41 902 ± 18 606 | 9,3E-2 |
| | 41B | $^{a}$36 349 ± 3 220 | $^{a}$36 294 ± 5 587 | $^{a}$37 936 ± 2 327 | $^{a}$38 281 ± 3 167 | 7,3E-1 |
| Fe | SO4 | $^{a}$1 595 ± 220 | $^{a}$1 363 ± 232 | $^{a}$1 782 ± 241 | $^{a}$1 486 ± 322 | 6,6E-2 |
| | 41B | $^{a}$2 033 ± 311 | $^{b}$1 290 ± 557 | $^{c}$2 945 ± 392 | $^{a}$2 017 ± 271 | 2,1E-5 |
| K | SO4 | $^{a}$28 548 ± 554 | $^{a}$27 171 ± 1 935 | $^{b}$22 647 ± 2 526 | $^{a}$27 425 ± 3 493 | 3,0E-3 |
| | 41B | $^{a}$31 393 ± 1 671 | $^{a}$28 832 ± 4 023 | $^{a}$30 989 ± 2 851 | $^{a}$29 173 ± 2 270 | 3,7E-1 |
| Mg | SO4 | $^{a}$5 303 ± 331 | $^{a}$5 102 ± 500 | $^{b}$3 952 ± 409 | $^{a}$5 170 ± 801 | 2,0E-3 |
| | 41B | $^{a}$5 509 ± 378 | $^{a}$5 713 ± 797 | $^{a}$5 331 ± 360 | $^{a}$5 222 ± 514 | 4,7E-1 |
| Mn | SO4 | $^{a}$196 ± 33 | $^{a}$130 ± 52 | $^{a}$147 ± 42 | $^{a}$188 ± 60 | 9,6E-2 |
| | 41B | $^{a}$226 ± 31 | $^{a}$211 ± 34 | $^{b}$289 ± 33 | $^{a}$212 ± 22 | 8,1E-4 |
| P | SO4 | $^{a}$1 793 ± 388 | $^{a}$1 595 ± 276 | $^{b}$1 137 ± 71 | $^{b}$1 059 ± 109 | 9,7E-05 |
| | 41B | $^{a}$2 195 ± 282 | $^{b}$1 869 ± 233 | $^{c}$1 394 ± 114 | $^{c}$1 346 ± 155 | 2,8E-06 |
| S | SO4 | $^{a}$3 191 ± 275 | $^{a}$3 062 ± 343 | $^{b}$2 573 ± 261 | $^{ab}$2 806 ± 165 | 5,0E-3 |
| | 41B | $^{a}$2 921 ± 162 | $^{a}$2 803 ± 264 | $^{a}$2 609 ± 161 | $^{a}$2 676 ± 302 | 1,6E-1 |
| Zn | SO4 | $^{a}$71 ± 9 | $^{a}$68 ± 10 | $^{a}$68.5 ± 8.1 | $^{a}$66.7 ± 5.2 | 3,0E-2 |
| | 41B | $^{a}$89 ± 7 | $^{b}$70 ± 16 | $^{ab}$76.1 ± 6.9 | $^{ab}$71.8 ± 6.3 | 8,6E-1 |

**Table S2.** Whole plant concentrations in ppm (mg/kg of DW) of Ca, Fe, K, Mg, Mn, P, S, Zn in SO4 and 41B under Ri, NoRi, WD and NoWD conditions. Values are mean ± sd, letters indicate significant differences according to adequate post-hoc comparisons.

| | | Leaves mineral content | | | | |
|---|---|---|---|---|---|---|
| | | Ri | | NoRi | | |
| Characteristic | | **NoWD** | **WD** | **NoWD** | **WD** | **P** |
| Ca | SO4 | $^{ab}$15 888 ± 614 | $^{a}$15 400 ± 1 486 | $^{a}$14 951 ± 4 083 | $^{b}$19 845 ± 2 767 | 1.8$^{E-2}$ |
| | 41B | $^{a}$17 446 ± 2 494 | $^{a}$20 288 ± 4 357 | $^{a}$20 695 ± 1 314 | $^{a}$19 366 ± 2 030 | 2,4$^{E-1}$ |
| Fe | SO4 | $^{a}$42.5 ± 6.1 | $^{a}$37.1 ± 2.5 | $^{a}$41.2 ± 6.8 | $^{a}$41.5 ± 7.9 | 4,8$^{E-1}$ |
| | 41B | $^{b}$43 ± 5 | $^{ab}$58 ± 7 | $^{a}$71 ± 13 | $^{b}$52 ± 10 | 8,6$^{E-4}$ |
| K | SO4 | $^{ab}$13 539 ± 1 942 | $^{a}$14 506 ± 2 167 | $^{b}$10 203 ± 2 495 | $^{a}$17 395 ± 3 420 | 1,0$^{E-3}$ |
| | 41B | $^{a}$11 380 ± 536 | $^{b}$15 300 ± 1 520 | $^{a}$10 440 ± 1 657 | $^{a}$11 123 ± 810 | 2,1$^{E-5}$ |
| Mg | SO4 | $^{ab}$2 286 ± 231 | $^{a}$2 452 ± 196 | $^{b}$1 761 ± 451 | $^{a}$2 790 ± 552 | 2,0$^{E-3}$ |
| | 41B | $^{a}$2 512 ± 150 | $^{a}$2 893 ± 575 | $^{a}$2 563 ± 341 | $^{a}$2 382 ± 437 | 2,4$^{E-1}$ |
| Mn | SO4 | $^{a}$98 ± 16 | $^{a}$70 ± 21 | $^{a}$81 ± 33 | $^{a}$87 ± 22 | 3,2$^{E-1}$ |
| | 41B | $^{a}$97 ± 14 | $^{a}$105 ± 14 | $^{a}$109 ± 17 | $^{a}$88 ± 6 | 6,4$^{E-2}$ |
| P | SO4 | $^{a}$740 ± 206 | $^{a}$633 ± 142 | $^{b}$427 ± 86 | $^{b}$393 ± 46 | 6,0$^{E-4}$ |
| | 41B | $^{a}$930 ± 150 | $^{a}$885 ± 102 | $^{b}$579 ± 102 | $^{b}$517 ± 83 | 6,3$^{E-6}$ |
| S | SO4 | $^{a}$1 361 ± 124 | $^{a}$1 343 ± 82 | $^{a}$1 146 ± 260 | $^{a}$1 274 ± 81 | 1,4$^{E-1}$ |
| | 41B | $^{ab}$1 355 ± 65 | $^{a}$1 438 ± 131 | $^{bc}$1 244 ± 81 | $^{c}$1 151 ± 107 | 8,0$^{E-4}$ |
| Zn | SO4 | $^{a}$28.81 ± 2.44 | $^{a}$27.19 ± 1.87 | $^{a}$27.42 ± 2.49 | $^{a}$28.69 ± 2.01 | 4,9$^{E-1}$ |
| | 41B | $^{a}$30.11 ± 1.92 | $^{a}$30.83 ± 3.83 | $^{a}$28.44 ± 2.58 | $^{a}$27.89 ± 3.52 | 3,7$^{E-1}$ |

**Table S3.** Leaf concentrations in ppm (mg/kg of DW) of Ca, Fe, K, Mg, Mn, P, S, Zn in SO4 and 41B under Ri, NoRi, WD and NoWD conditions. Values are mean ± sd, letters indicate significant differences according to adequate post-hoc comparisons.

| Root mineral content | | | | | | |
|---|---|---|---|---|---|---|
| | | **Ri** | | **NoRi** | | |
| Characteristic | | **NoWD** | **WD** | **NoWD** | **WD** | **P** |
| Ca | SO4 | [a]15 425 ± 1 743 | [a]16 421 ± 2 813 | [a]13 955 ± 1 238 | [a]22 057 ± 18 099 | $2,3^{E-1}$ |
| | 41B | [a]18 903 ± 1 222 | [a]16 006 ± 3 757 | [a]17 241 ± 1 418 | [a]18 915 ± 1 895 | $1,4^{E-1}$ |
| Fe | SO4 | [a]1 553 ± 218 | [a]1 326 ± 231 | [a]1 741 ± 245 | [a]1 444 ± 320 | $6,9^{E-2}$ |
| | 41B | [b]1 990 ± 310 | [c]1 232 ± 559 | [a]2 874 ± 391 | [b]1 965 ± 271 | $2,3^{E-5}$ |
| K | SO4 | [a]15 008 ± 2 146 | [ab]12 665 ± 1 930 | [ab]12 444 ± 1 445 | [b]10 031 ± 1 396 | $2,0^{E-3}$ |
| | 41B | [a]20 013 ± 1 768 | [b]13 532 ± 4 253 | [a]20 549 ± 2 509 | [ab]18 050 ± 2 056 | $3,0^{E-3}$ |
| Mg | SO4 | [a]3 016 ± 224 | [ab]2 650 ± 393 | [b]2 191 ± 265 | [b]2 380 ± 434 | $5,0^{E-3}$ |
| | 41B | [a]2 997 ± 315 | [a]2 820 ± 568 | [a]2 768 ± 187 | [a]2 839 ± 307 | $7,6^{E-1}$ |
| Mn | SO4 | [a]98 ± 22 | [a]59 ± 31 | [a]66 ± 15 | [a]101 ± 41 | $5,3^{E-2}$ |
| | 41B | [b]128 ± 24 | [b]106 ± 29 | [a]181 ± 28 | [b]124 ± 21 | $7,6^{E-4}$ |
| P | SO4 | [a]1 054 ± 189 | [a]962 ± 137 | [b]710 ± 54 | [b]666 ± 110 | $1,1^{E-4}$ |
| | 41B | [a]1 265 ± 187 | [b]984 ± 192 | [b]815 ± 73 | [b]830 ± 111 | $2,3^{E-4}$ |
| S | SO4 | [a]1 830 ± 251 | [ab]1 719 ± 304 | [b]1 428 ± 148 | [ab]1 533 ± 204 | $5,0^{E-2}$ |
| | 41B | [a]1 566 ± 120 | [a]1 365 ± 231 | [a]1 365 ± 105 | [a]1 524 ± 250 | $2,0^{E-1}$ |
| Zn | SO4 | [a]42 ± 8 | [a]41 ± 10 | [a]41.1 ± 7.9 | [a]38.0 ± 5.1 | $8,5^{E-1}$ |
| | 41B | [a]59 ± 8 | [b]39 ± 13 | [ab]48 ± 8 | [ab]44 ± 7 | $2,9^{E-2}$ |

**Table S4.** Root concentrations in ppm (mg/kg of DW) of Ca, Fe, K, Mg, Mn, P, S, Zn in SO4 and 41B under Ri, NoRi, WD and NoWD conditions. Values are mean ± sd, letters indicate significant differences according to adequate post-hoc comparisons.

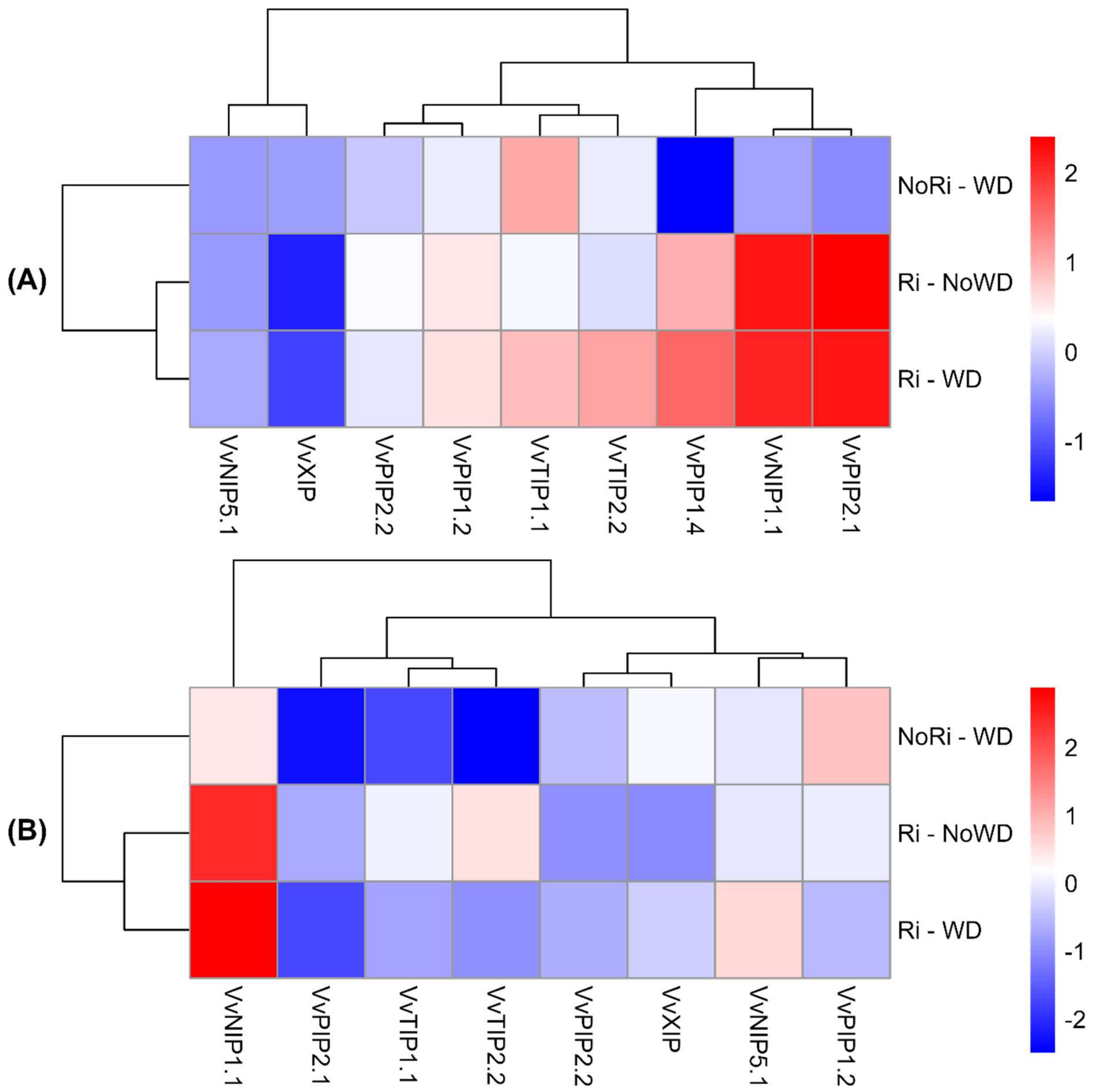


**Fig S1. Effect of Ri symbiosis on the expression of aquaporin genes in rootstock roots under NoWD and WD regimes.**

Heatmaps represent the expression variation compared to NoRi – NoWD condition for 41B (A) and SO4 (B).
